# Establishment of Wnt ligand-receptor organization and cell polarity in the *C. elegans* embryo

**DOI:** 10.1101/2023.01.17.524363

**Authors:** Pierre Recouvreux, Pritha Pai, Rémy Torro, Mónika Ludányi, Pauline Mélénec, Mariem Boughzala, Vincent Bertrand, Pierre-François Lenne

## Abstract

Different signaling mechanisms concur to ensure robust tissue patterning and cell fate instruction during animal development. Most of these mechanisms rely on signaling proteins that are produced, transported and detected. The spatiotemporal dynamics of signaling molecules is largely unknown, yet it determines signal activity’s range and time frame. Here, we use the *Caenorhabditis elegans* embryo to study how Wnt ligands, an evolutionarily conserved family of signaling proteins, dynamically organize to establish cell polarity in a developing tissue. We identify how locally produced Wnt ligands spread to transmit information to distant target cells. With quantitative live imaging, we show that the Wnt ligands diffuse extracellularly through the embryo over a timescale shorter than the cell cycle. We extract diffusion coefficients of Wnt ligands and their receptor Frizzled (Fz) and characterize their co-localization. Integrating our different measurements and observations in a simple computational framework, we show how fast diffusion in the embryo can polarize target cells. Our results support diffusion-based long-range Wnt signaling, which is consistent with the dynamics of developing processes.

## Introduction

Animal development results from morphogenetic processes that must be spatially and temporally coordinated to pattern tissues, instruct cell fates, or orient cell divisions. Coordination originates from interactions between cells or groups of cells either in contact or at a distance. Intercellular transfer of information is achieved by signaling molecules whose role is to activate signaling pathways or polarize cells. Morphogens are such signaling molecules that spread in tissues and typically form concentration gradients which are transduced in signaling gradients, leading to tissue patterning. Even if signaling gradients are very often considered to be at play in multiple organisms, only a few studies report the observation of signaling protein concentration gradients *in vivo* ^1–5^. Following the movement of signaling molecules in live tissue is challenging. Therefore, to a few exceptions, the modes of dispersion of morphogens in tissues remain unclear. It is often proposed that morphogens diffuse in the extracellular space and establish a concentration gradient, whose characteristic length could be set by the stability of the protein, its diffusion properties, its interactions with the extracellular matrix, or its receptors ^6,7^. It is essential to better understand the mode of dispersal of signaling molecules in tissues because it imposes spatial and temporal constraints on the timing and range at which signaling can occur. In particular, one can hypothesize that a fast-developing tissue necessitates rapid dispersal modes, while a slowly-growing tissue can rely on dispersal with slower dynamics. For molecular transfer of signals from source to target cells, organisms have developed mechanisms adapted to their developmental dynamics and sizes, which impose constraints on the possible processes of signaling molecule spreading. Over short distances, typically a few cell diameters, diffusion can establish concentration gradients, while over long distances, it is unclear how such a process can establish a robust signaling gradient in a suitable timeframe. It is often proposed that free diffusion is complemented or replaced by active transport mechanisms such as transcytosis or cytonemes in order to increase the robustness or the range of signaling ^8^. Until now, free diffusion of signaling molecules has only rarely been reported in living tissues during development ^5,9^. Here, we analyzed the existence of a freely diffusing population of Wnt ligands in *C. elegans* embryos and its potential ability to instruct cell fate decisions.

Wnt is a family of conserved signaling proteins that play crucial roles during development and homeostasis in metazoans. Wnt signaling activity is generally graded along the primary body axis over short or long distances ^10^. Various mechanisms have been proposed to explain these ranges. Though Wnt is often believed to be a secreted protein that signals at a distance as a diffusible molecule in a developing tissue, evidence for that has been only reported in a few living tissues ^4,11^. An important point to consider is the timescales over which Wnt signaling must be established. Depending on the different systems considered, this timescale can range from tens of minutes to tens of hours; it thus imposes constraints on the mode of dispersion of the ligand, which is also dependent on the range over which signaling must occur ^12^.

In *C. elegans*, Wnt signaling has been shown to convey polarity information to target cells along the anteroposterior axis by controlling the orientation and asymmetry of divisions ^13–15^. These events instruct fate decisions. It is important to consider that such polarizing mechanisms must occur in a timeframe consistent with the cell cycle, which is, in the case of *C. elegans* embryo, about ½ hour. To better understand the role of Wnt ligands in the polarization of *C. elegans* embryos, we took advantage of the recent possibility to fluorescently tag Wnts in living *C. elegans* ^4,16–18^. We characterized the distribution of Wnt ligands and their receptor Fz and evidenced the existence of a freely diffusing population of the ligand. We further analyzed the interplay between the ligand and the receptor in live embryos and in single embryonic cells culture. Integrating our data on the dynamics of Wnt ligands and their receptor in a simple physical model, we computationally explore how cell-scale polarity can be established. Our simulations indicate that a free diffusion-based mechanism could convey polarity information in a developing embryo over a timeframe shorter than the cell cycle.

## Results

### Wnt ligands are produced posteriorly and form clusters along the anteroposterior axis

*C. elegans* has five Wnt ligands ^19^ which are not all expressed simultaneously during embryogenesis ^18,20,21^. *cwn-1, cwn-2*, and *mom-2* are expressed in embryos before elongation, while *egl-20* and *lin-44* expressions start at later stages. To study the spatiotemporal dynamics of these Wnt ligands before elongation, we labeled CWN-1, CWN-2, and MOM-2 proteins with YFP ^18^. The three Wnt ligands are detected in live embryos at different stages with a similar distribution [Figure 1b-d]. They appear either as membrane-bound punctae [Figure 1a cyan arrowheads] or as a more diffused signal in the cytoplasm of cells [Figure 1a magenta arrowheads]. Cells with a cytoplasmic signal are located in the posterior half of embryos; they are the source of Wnt, as also suggested by earlier observations on transcriptional reporters or mRNA that showed a graded transcription of *cwn-1, cwn-2* and *mom-2* at the posterior end ^18,20,21^. While the three Wnt ligands display similar distributions, there are differences in their sources’ locations, sizes and expression levels. CWN-2::YFP has the largest source, as shown by the extent of cells with a cytoplasmic signal [Figure 1b]. CWN-1 [Figure 1c] and MOM-2 [Figure 1d] sources are more restricted spatially and located closer to the posterior tip. For the three ligands, we observed an increase in total fluorescence signals over time; CWN-2::YFP is the most precocious and shows the strongest signal, while CWN-1::YFP and MOM-2::YFP are detected later and with lower intensities [Figure 1e & Figure 1 – supplementary figure 1a]. The total concentration of CWN-2::YFP varies along the anteroposterior axis [Figure 1g], with a maximum in the source region where the cytoplasmic signal dominates. This polarity at the tissue level increases over time and is maximal at about 4 hours, and the total signal is then twice higher in the posterior half than in the anterior half of the embryo [Figure 1f]. Although CWN-1::YFP, CWN-2::YFP, and MOM-2::YFP are produced in the background of endogenous Wnt ligands, we did not observe any phenotype associated with potential overexpression.

**Figure 1.**
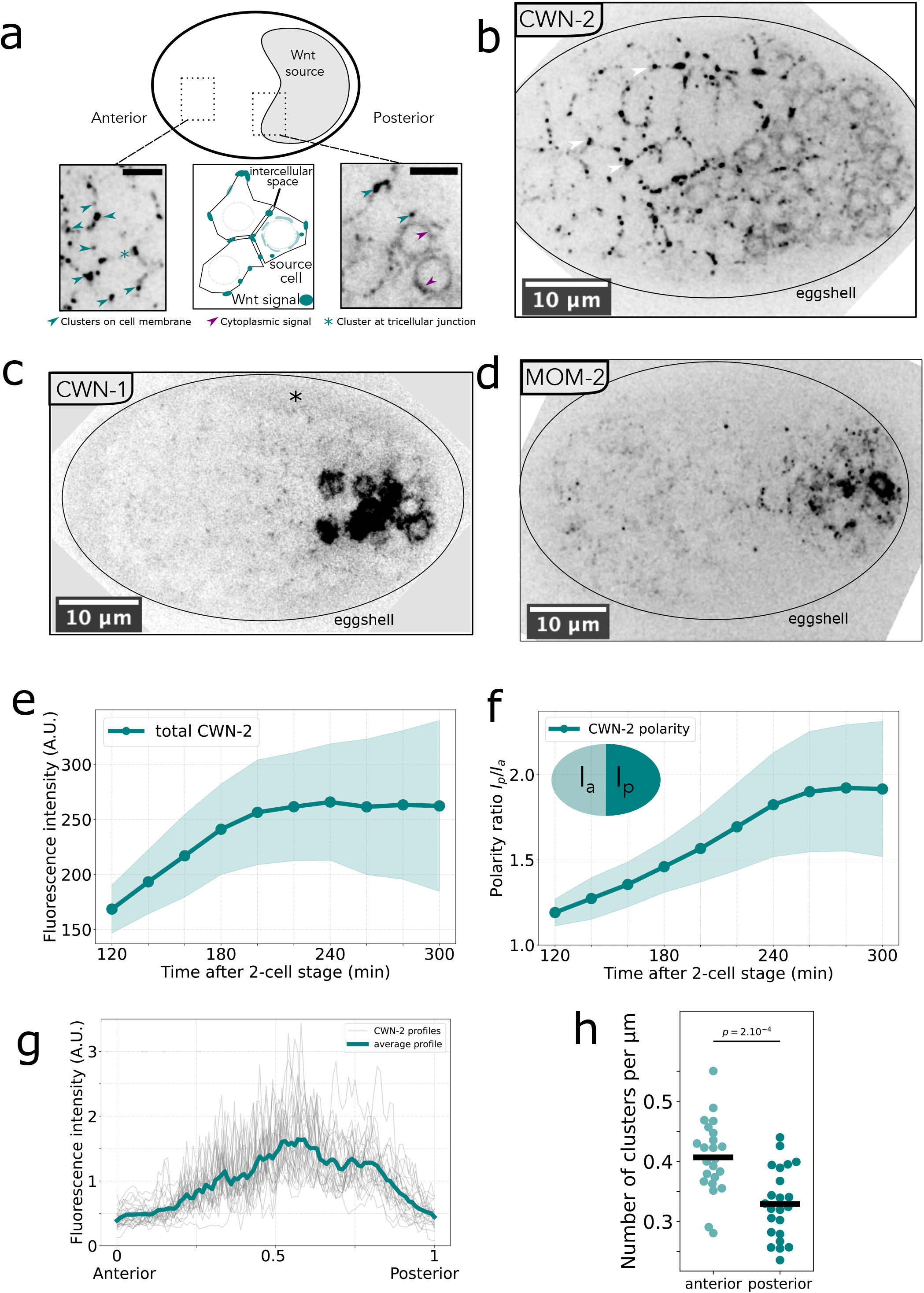
Distribution of Wnt ligands in *C. elegans* embryos. a- Schematic representation of the position of Wnt source cells (top) and close up on CWN-2::YFP localization at the cell level. The signal is found in the cytoplasm of source cells (magenta arrowheads) and as clusters on the cell membrane (cyan arrowheads). Some of the puncta are detected at tricellular junctions (cyan asterisk). Scale bar 4µm. b-, c-, d- Fluorescence microscopy images of 4 hour-old embryo expressing CWN-2::YFP (*vbaIs43*), CWN-1::YFP (*vbaIs39*), MOM-2::YFP (*vbaIs40*). White arrowheads in b indicate clusters at tricellular junctions. The three ligands show a similar pattern: source cells are located at the posterior end of the embryo and puncta are detected on the cell membranes. A diffuse fluorescence signal is observed between the embryonic tissue and the eggshell (asterisk panel c). e- Time evolution of total fluorescence intensity of CWN-2::YFP (*vbaIs43*). Acquisition starts 2h after the 2-cell stage. f- Time evolution of the polarity ratio. Defined as the ratio of the total fluorescence signal in the posterior half to the total signal in the anterior half. g- Antero-posterior profile of CWN-2::YFP (*vbaIs43*) fluorescence intensity. In gray single embryo profiles, in green average profile (n=26 embryos). h- Number of CWN-2::YFP (*vbaIs43*) clusters on cell membranes in the anterior and posterior halves (n= 23 embryos).

Next, we addressed the localization of the membrane-bound clusters. They are detected anteriorly, away from the sources from which Wnt proteins are likely to originate. Previous studies have shown similar punctae on membranes for another Wnt ligand EGL-20 ^1,4^. We observed that CWN-2::YFP clusters were moving with the membrane though no noticeable movement along the membrane was detected [movie 1]. However, the movement of cell-cell contacts due to tissue reorganization makes it challenging to track in 3D these objects over timescales longer than a minute. Notably, these Wnt clusters were often detected at tricellular junctions [Figure 1a-b, movie 1]. Importantly, we noticed that punctae density is lower on the membrane of CWN-2 source cells than on the membrane of cells with no cytoplasmic signal in the anterior half of the embryo [Figure 1 b & h], suggesting that the clustering of Wnt ligands at the membrane is regulated.

### Wnt ligands diffuse in the extracellular space

The presence of Wnt ligands in the anterior region of the embryo, while their source cells are more posterior, raises the possibility that Wnt ligands move over long distances. We thus hypothesized that Wnt ligands could diffuse in the extracellular space, between the cells and between the embryonic tissue and the eggshell [asterisk in Figure 1c]. To directly test this hypothesis, we performed Fluorescence Correlation Spectroscopy (FCS) experiments on MOM-2::YFP in that space [Figure 2a]. Fluorescence intensity autocorrelation curves show that the detected species freely diffuse with a diffusion coefficient of about 10 µm^2^/s [Figure 2b], consistent with a protein of the size and nature of Wnt ^11,22^. The localization and the existence of a secreted Wnt population outside the embryonic tissue support the idea that Wnt clusters are present in the extracellular space and originate from the local clustering of secreted ligands diffusing in the extracellular space. To further prove that Wnt ligands can diffuse in the extracellular space within the tissue and contribute to their characteristic distribution, we performed FRAP (Fluorescence Recovery After Photobleaching) experiments [Figure 2d and movie 2]. We monitored the recovery of the CWN-2::YFP signal in the bleached anterior half of the embryo (n=6). We observed the reappearance of punctae within the first minute [Figure 2c red] in the anterior compartment closest to the source [red box in Figure 2d]. In comparison, the recovery in the most anterior compartment [blue box in Figure 2d] is delayed by a few minutes [Figure 2c blue]. We estimate the characteristic time of clusters recovery to be about 200s [Figure 2c dashed line]. This result indicates that free Wnt ligands originating from the posterior half diffuse between cells, reach the anterior part of the embryo, and accumulate in clusters. Interestingly, for a species with a diffusion coefficient of 10µm^2^/s, the timescale for diffusion over a length scale of 20 µm (half the size of an embryo) is below one minute. We thus measure a recovery presumably driven by molecules with a slower diffusion coefficient. This discrepancy could be explained by the fact that Wnt ligands have a lower effective diffusion coefficient in the intercellular space because of possible interactions with the extracellular matrix, plasma membranes, and its receptors. Our results confirm the existence of a population of Wnt ligands that can spread over long distances in a timescale consistent with free diffusion and significantly shorter than the duration of the cell cycle. To better understand the formation of such punctae in the tissue, we decided to study the distribution of a Wnt receptor of the Frizzled family, MOM-5, which is essential for *C. elegans* embryogenesis and polarity ^21,23^.

**Figure 2.**
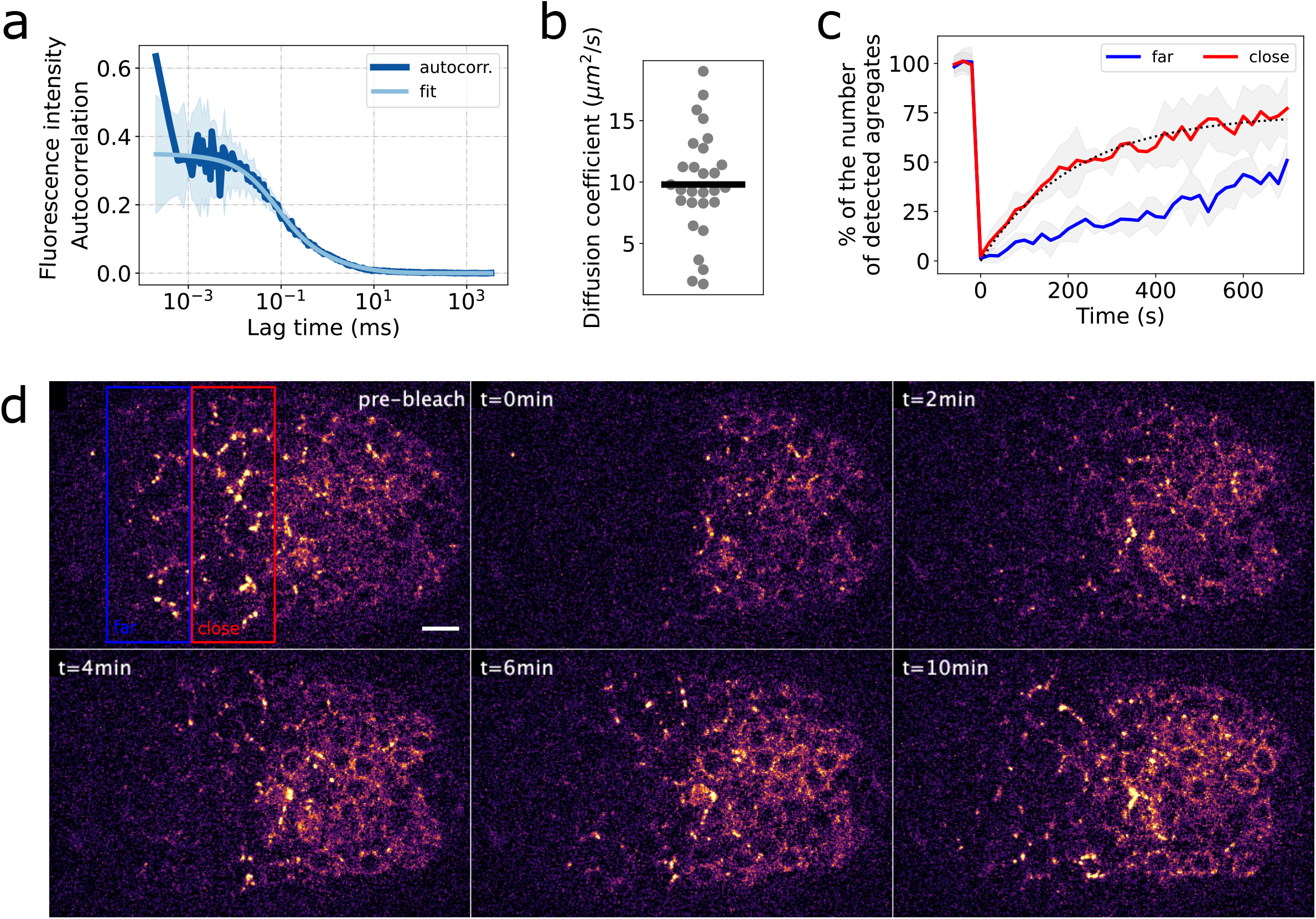
Dynamics of Wnt in embryos. a- Average (+/-std) fluorescence intensity autocorrelogram (dark blue) from FCS experiment on MOM-2::YFP (*vbaIs40*) and fitting of the average (light blue) (n=28, 3 embryos). Measurement was done in between the embryonic tissue and the eggshell (see Fig 1c). b- Diffusion coefficients of MOM-2::YFP outside the embryonic tissue estimated from fits of individual autocorrelograms. In that space, the molecule has an average diffusion constant of 10µm^2^/s. c- Recovery of the number of detected clusters after photobleaching (see movie 2). At time 0 the anterior half of the embryo is bleached, and recovery is measured in two distinct zones (i) *close* corresponds to the red box in d and (ii) *far* corresponds to the blue box in d. The recovery curve in *close* is fitted by an exponential (dotted line) that gives a characteristic of about 200s. d- Representative images from movie 2 of the FRAP experiment showing the bleached zone and recovery over time. Scale bar 5 µm.

### Fz/MOM-5 distribution is not uniform

We used strains with endogenous protein tagged with fluorescent proteins to study the spatiotemporal distribution of Wnt receptor Frizzled/MOM-5 on the plasma membrane [Figure 3a]. We measured the time evolution of total endogenous Fz/MOM-5 (MOM-5::YPET *cp31*) ^17^) and observed that the signal is detectable at early stages and increases over time [Figure 3b]. At the tissue level, the signal is not homogeneous along the anteroposterior axis. The ratio posterior-vs-anterior signals for MOM-5::YPET decreased over time [Figure 3c]. Interestingly, the posterior half of the embryo, where the Fz/MOM-5 signal is lower at 4 hours after the 2-cell stage, is the region in which Wnt source cells are located and where the Wnt signal is highest [Figure 3d]. Fz/MOM-5 and Wnt show opposite localizations at the tissue scale, MOM-5 being lowest where Wnt levels are highest [Figure 2e and 3c].

**Figure 3.**
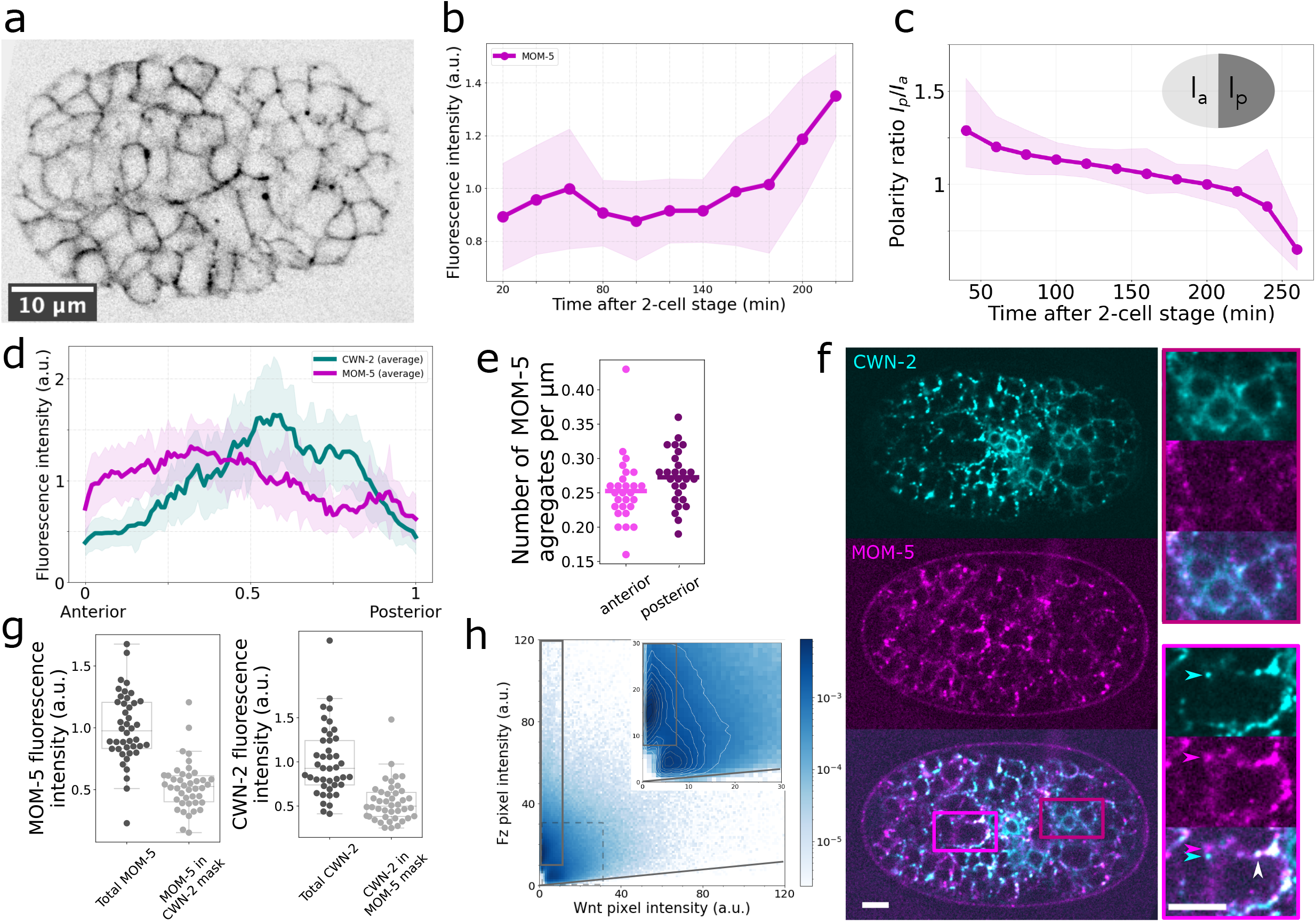
Fz/MOM-5 and Wnt/CWN-2 distributions in embryos. a- Fluorescence confocal section showing the endogenous MOM-5 tagged with YPET (*cp31*) at 4 hours after the 2-cell stage. b- Time evolution of the total fluorescence intensity of MOM-5::YPET. MOM-5::YPET is detectable early on and its level increases around 2.5 hours after the 2-cell stage (n=6 embryos). c- Time evolution of the polarity ratio defined as the membrane fluorescence intensity of MOM- 5::YPET in the posterior half divided by the one in the anterior half. As from 3 hours after 2-cell stage MOM-5::YPET signal in the posterior half of the embryo is lower than in the anterior half (n=6 embryos). d- Fluorescence intensity anteroposterior profiles at 4 hours after the 2-cell stage for CWN-2::YFP (*vbaIs43*) (in green) and MOM-5::YPET (*cp31*) (in magenta). e- Number of MOM-5::mKate2 (*vba10*) puncta per µm of the apposed membranes in the anterior and posterior halves (n=10 embryos). f- Confocal fluorescence slices of an embryo expressing MOM-5::mKate2 (magenta) and CWN-2::YFP (cyan). CWN-2 and MOM-5 are detectable separately (cyan and magenta arrowheads in zoom) or together (white arrowhead in zoom). Scale bar = 5 µm. g & h – Analysis of the fluorescence intensity of CWN-2::YFP and MOM-5::mKate2 in thresholded images g- (left panel) MOM-5::mKate2 total membrane fluorescence intensity (left) and intensity when CWN-2::YFP is detected in the same pixel (right). (right panel) CWN-2::YFP total membrane fluorescence intensity (left) and intensity when MOM-5::mKate2 is detected in the same pixel (n=41, 15 embryos). h- 2D-histogram of the Fz and Wnt fluorescence intensities in the same pixels. Colorbar indicates the frequency of occurrence (n=15 embryos).

At the cellular level, Fz/MOM-5 is not homogeneously distributed on the plasma membrane and forms clusters of higher intensity on top of a low uniform signal [Figure 3a and 3f]. We estimated the number of clusters per µm of the plasma membrane in embryos to about 2 to 3 clusters per 10 µm of apposed membranes [Figure 3e] (n=27 in 10 embryos). Even though the MOM-5::YPET signal is lower in the posterior half compared to the anterior one, cluster density is similar in both compartments [Figure 3e].

### MOM-5::mKate2 and CWN-2::YFP partially co-localize on the plasma membrane

It has been previously shown that the Wnt ligand’s spatial distribution can affect the localization of their receptors in various systems ^18,24,25^. In *C. elegans* Fz/MOM-5 distribution is transiently enriched at the posterior poles of many dividing cells during embryogenesis ^18,23^. Therefore we decided to investigate the relative localization of CWN-2 and MOM-5 at the cellular level to see if high or low signals in one of the two species correlate with the signal amplitude of the other. We prepared a strain in which the coding sequence of mKate2 fluorescent protein was inserted at the C-terminus of the Fz/MOM-5 endogenous locus (see Mat & Meth, MOM-5::mKate2 (*vba10*)) and imaged CWN-2::YFP and MOM-5::mKate2 in the same embryos [Figure 3f]. We observed that Fz punctae can be found with [white arrowhead figure 3f] or without CWN-2 clusters [magenta arrowhead Figure 3f], and CWN-2::YFP clusters can also be found in the absence of a MOM-5:mKate2 cluster [cyan arrowhead Figure 3f]. We estimated that about half of the total quantity of MOM-5::mKate2 (resp. CWN-2::YFP) is found in pixels that coincide with detectable CWN-2::YFP (resp. MOM-5::mKate2) clusters [Figure 3g] (n=42 in 21 embryos). This observation points to their local interaction but also raises the possibility that the organization Wnt ligands/receptors are only partially dependent. We thus tested the influence of the local concentration of the ligand (resp. receptor) on the concentration of the receptor (resp. ligand) by extracting the same-pixel intensities of MOM-5::mKate2 and CWN-2::YFP [Figure 3f], in 4-hour-old embryos (n=42). For example, one could expect that high local levels of the ligand could recruit the receptor and correlate with high receptor levels. Reciprocally, receptor clusters could act as platforms for the recruitment of the ligands. We plotted the ligand/receptor intensities 2D-histogram [Figure 3h] to study this possible correlation. We could observe that when the ligand concentration increases, the receptor concentration also tends to increase (gray line Figure 3h), supporting the idea that high ligand levels correlate with higher levels of the receptor. However, we also observed that high levels of the receptors could be detected with low levels of the ligands (gray rectangle figure 3h). It suggests that the clustering of Fz/MOM-5 is at least partially independent of CWN-2 ligands. The rest of the distribution shows that the relationship between Fz and Wnt levels is complex and not based on direct correlations. This result is also supported by other experiments in which we look at the distribution of CWN-2::YFP in the absence of Fz/MOM-5 [Movie 3]. Even though the tissue was disturbed, CWN-2 was expressed in source cells, and we detected clusters on cellular membranes, supporting the idea that Wnt puncta can form, at least in part, independently of Fz/MOM-5.

### MOM-5/Fz dynamics on the plasma membrane of dissociated cultured embryonic cells

The distribution of Fz receptors on the plasma membranes results from the balance between transport toward the membrane, dynamics in the membrane, and recycling to the cytoplasm. We wanted to characterize the quantity and the dynamics of receptors on the plasma membrane to understand the formation of Fz clusters. We complemented our *in vivo* spatial characterization of clusters with an *in vitro* assay that circumvent some limitations of the *in vivo* approach. Measuring the dynamics of receptors *in vivo* is highly challenging as cell-cell contacts are constantly moving and individual plasma membranes of neighboring cells cannot be resolved optically. Thus we decided to study MOM-5::YPET distribution and dynamics in dissociated single embryonic cells in culture. After egg collection and eggshell digestion, embryonic tissues were mechanically dissociated and centrifuged to obtain single isolated *C*.*elegans* embryonic cells [Figure 4a]. We measured the plasma membrane distribution of MOM-5::YPET in cultures (*cp31*) and found that the receptor, compared to a membrane marker, was not homogeneously spread on the membrane [Figure 4b]. We counted, on average, about 1 cluster per 10µm of the plasma membrane [Figure 4c], in the same range as what is obtained in embryos for a single membrane [Figure 3e]. These results suggest that the cluster organization of the receptor on the membrane exists independently from direct cell-cell contacts and might result from direct or indirect interactions within the membranes or in the cytoplasm. We then measured the dynamics of receptors by tracking single MOM-5::YPET molecules at a high framerate (50 ms) [movie 4]. We obtained the trajectories of these molecules with the ImageJ plugin TrackMate ^26^ and analyzed their mean square displacements (MSD). We fitted MSDs [Figure 4d] with 4Dt*α*, with D the diffusion coefficient and *α* the anomalous diffusion exponent ^27^ that characterize the type of diffusion (*α* ∼ 1 for free, *α* <1 for confined and *α* >1 for directed diffusions). We calculated the diffusion coefficient of Frizzled (D) from diffusive trajectories (0.9 < *α* < 1, n=220). We found an average of 0.19μm^2^/s [Figure 4 c-d], a value typical of a membrane receptor.

**Figure 4.**
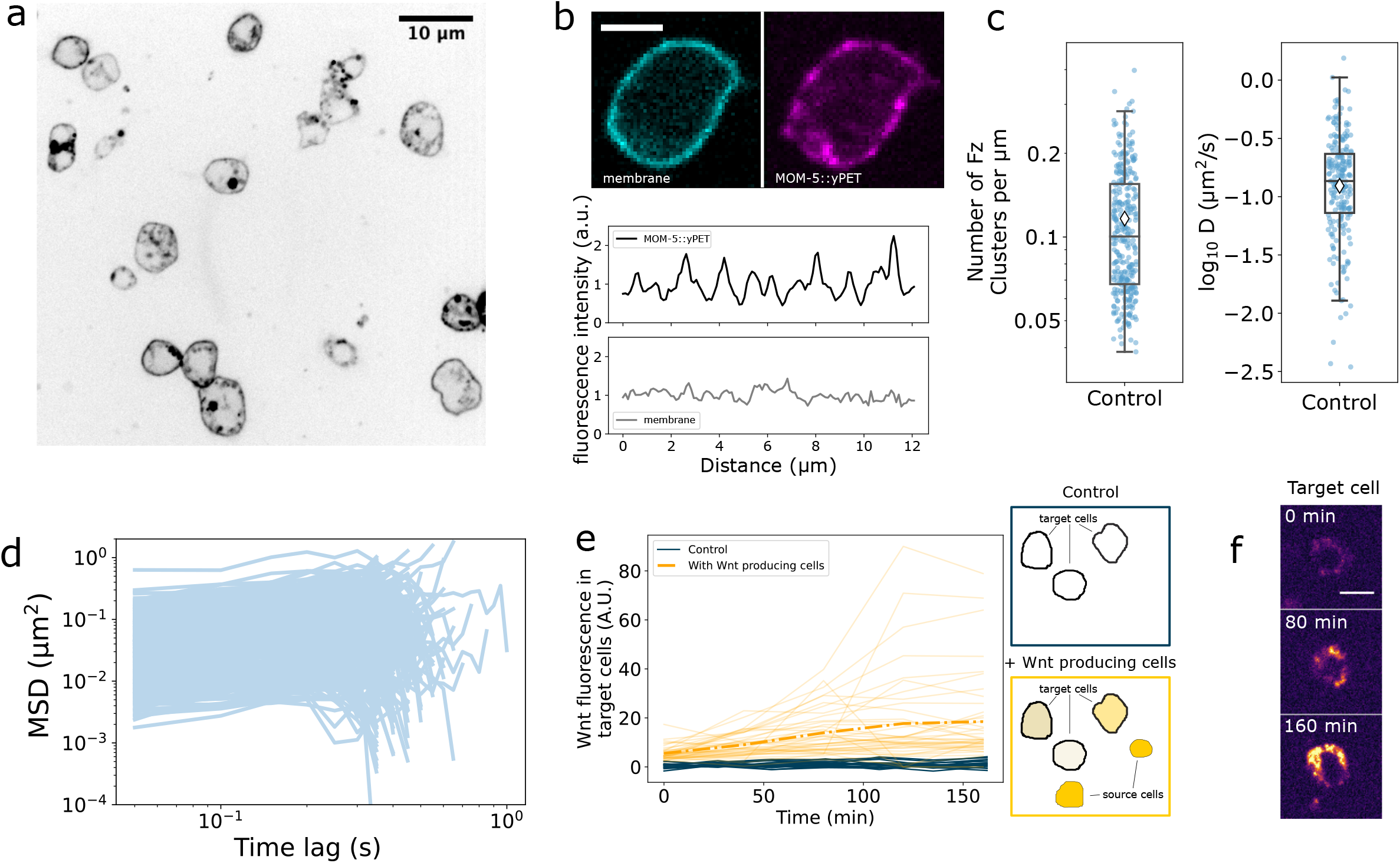
Fz distribution and dynamics on the membrane of embryonic dissociated cells. a- Culture of dissociated embryonic cells expressing MOM-5::YPET (*cp31*). b- (top) Fluorescence images of a cell expressing a membrane marker (*ItIs44*, mCherry::PH) (cyan) and MOM-5::YPET (magenta); (bottom) Normalized (mean) fluorescence intensity profiles along the plasma membrane of the membrane marker and of MOM-5::YPET, showing the more heterogenous distribution of Fz. c- *left* Number of Fz/MOM-5::YPET clusters per µm along the plasma membrane of single embryonic cells. *right* Diffusion coefficients of MOM-5/Fz in dissociated cells, obtained from data presented in d. d- Mean Square Displacement (MSD) curves obtained from Fz/MOM-5::YPET trajectories. e- Time evolution of the cytoplasmic fluorescence signal of CWN-2::mCherry (*vbaIs29*) in MOM-5::YPET expressing cells in the presence (yellow) or absence (blue) of CWN-2::mCherry expressing cells. *right* cartoon of the experimental cell condition in control (top, MOM-5::YPET expressing cells in absence of Wnt producing cells) and (bottom) in the presence of CWN-2::mCherry expressing cells. f- Snapshots of CWN-2::mCherry fluorescence signal in MOM-5::YPET expressing cells. Scale bar is 4µm.

We also co-cultured single isolated embryonic cells in the presence of *C. elegans* isolated cells from another strain that overexpresses a mCherry tagged version of CWN-2 (*vbaIs29*). After heat shock, these cells express CWN-2::mCherry. Interestingly, we could observe a gradual accumulation of the fluorescent mCherry signal in the cytoplasm of MOM-5::YPET expressing cells [Figure 4e-f]. These results can be explained by the secretion of CWN-2::mCherry by producing cells. Secreted ligands reaching MOM-5::YPET cells are trapped and transported into the cytoplasm, where it accumulates. This observation supports that Wnt ligands can diffuse freely in the extracellular space, even without potential interactors of the extracellular matrix.

### Polarity transfer from tissue to cellular scale

We have observed that Wnt ligands are produced in the posterior half of the embryo and diffuse toward the anterior half with a characteristic time of a few minutes. It is important to highlight that this diffusion is rapid compared to the cell cycle (20 to 30 minutes). A fundamental question is how this polarity at the tissue level is transferred to the cellular level. Indeed, it is known that many cells in the embryo can distinguish anterior from posterior and take subsequent fate decisions based on Wnt signaling ^13–15^. It has even been suggested that Wnt ligands could polarize its receptor Fz in *C. elegans* ^18,24^. These cells must measure this information on a timescale shorter than the cell cycle. To address this question, we performed numerical simulations of the diffusion of the ligand and its interaction with the receptor in a 2D idealized embryo(software Smoldyn)^28,29^. We placed 12 columns and 8 rows of identical square cells (4µm) separated by 50 nm [Figure 5a-b]. The posterior column (#12, rightmost column) secretes Wnt ligands that will then diffuse in between cells. On the membrane of these cells, we placed Frizzled receptors, able to diffuse in the membrane. We chose the diffusion coefficients of these species based on our experimental results (D_Wnt_ = 2 µm^2^/s and D_Fz_ = 0.1 µm^2^/s). Fz and Wnt can bind together and form a complex through a diffusion-limited reaction. We considered the complex formed to be immobile on the membrane, to facilitate the localization of ligand-receptor interaction events, upon which downstream signaling events (not simulated) can be established. We defined a cell polarity index 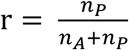, with *n*_*p*_ (resp. *n*_*A*_) the number of complexes in the posterior (resp. anterior) half of a cell [Figure 5c]. We observed [movie 5] the progression toward the anterior of a front of diffusing Wnt while diffusing the ligands bind to the receptors. Interestingly, we found that it was more likely to form complexes on the posterior side of the cells than on the anterior side (*r* % > 0.5). We observed that r is higher for cells further away from the source; on the opposite, cells close to the source of Wnt showed almost no polarity (%~0.5). In a given column of cells, the average polarity index is time-dependent [Figure 5b] and tends to decrease toward the steady-state value [Figure 5b red line]. This can be explained by the fact that the Wnt wave hits first the posterior side, creating complexes, while the anterior half has not yet been exposed to ligands. Nevertheless, the steady-state polarity index of cells further away from the source is significantly larger than 0.5. This indicates that the polarity established at the tissue level can be transmitted to the cellular level at a distance from the source. However, if the ligand diffuses faster (10 µm^2^/s), we observe that the polarity of a given cell column is decreased [Figure 5c]; only cells further from the source have a polarity index over 0.5. In this condition, a significant fraction of cells in the embryo is not polarized. In the other limit, if Wnt is diffusing slower (0.1 µm^2^/s) the polarity index will increase. Still, then the ligand will need a much longer time to reach cells in the anterior part, impeding the transfer of polarity over a large scale [Figure 5c] in the timescale of the cell cycle.

**Figure 5.**
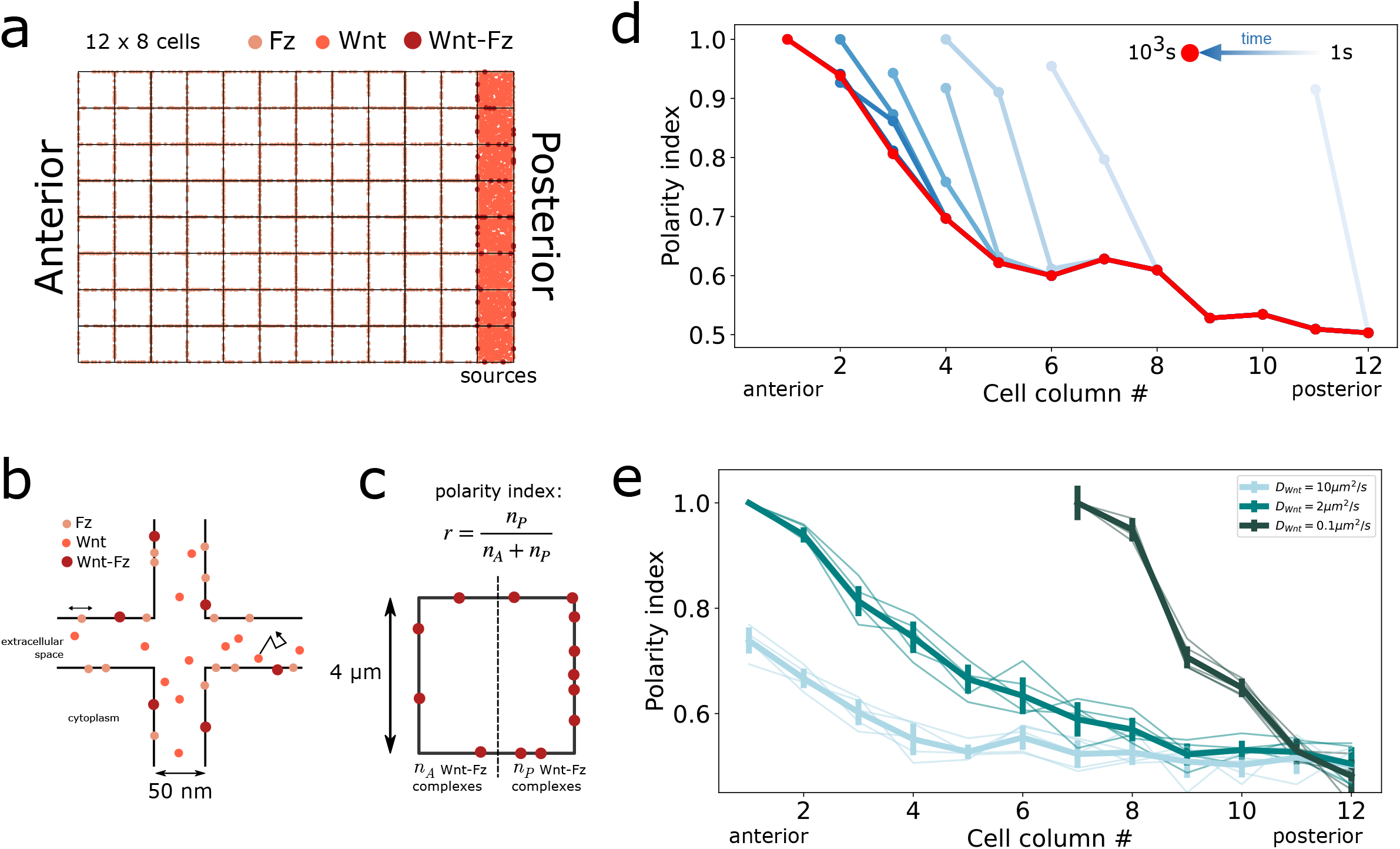
Simulations of the transfer of polarity from the tissue scale to the cell scale. a & b- Geometry of simulations. 96 square cells of 4 µm are placed on 8 rows and 12 columns. Cells are separated by 50 nm. Source cells (posterior column) express and secrete Wnt in extracellular space. Fz molecules are placed on the plasma membrane and diffuse in the membrane (D =0.1µm^2^/s), Wnt molecules are secreted from the rightmost columns (posterior) and diffuse in between cells (D_Wnt_ = 2 µm^2^/s). Wnt-Fz complexes are formed on membranes upon binding between Wnt and Fz molecules. Wnt-Fz do not diffuse, their location corresponds to where they are formed. c- The cell polarity index is defined as the ratio between the number of complexes in the posterior half (n_P_) and the total number of clusters (n_A_+n_P_). d- Average polarity index of the different cell columns over time, from 0s (light blue) to 10^3^s (red). The polarity index is not defined in every column at early time points because Wnt has not yet reached the anterior columns. e- Polarity index in columns at t=10^3^s for different Wnt diffusion coefficients for 5 simulations per condition. Average and standard deviation in bold.

## Discussion

In this study, we described the distribution of Wnt ligands and their receptors, MOM-5/Frizzled, in *C. elegans* embryos and at the surface of cultured embryonic cells, to determine the mechanisms at play in Wnt spreading and in the regulation of Wnt signaling activity. We showed that Wnt ligands (CWN-1, CWN-2 and MOM-2) are produced in the posterior half of embryos and are secreted in the extracellular space. They form stable punctae on the plasma membrane of cells in the region of the source as well as away from this source [Figure 1]. This distribution is similar to that found in *C. elegans* at later stages and for a different Wnt ligand ^1,4^. We have shown that these structures can form independently of the receptor MOM-5/Frizzled [Figure 3 and Movie 3], suggesting a possible role for other membrane proteins (such as other transmembrane receptors) or extracellular components which are known to affect Wnt activity like Heparan-Sulfate Proteoglycans (HSPGs) or secreted Frizzled-Related Proteins (sFRP) ^11,20,30,31^. Notably, we evidenced that besides these clusters, a population of Wnt ligands is spreading in the tissue with dynamics consistent with free diffusion of the protein of the size of Wnt [Figure 2]. However, due to the hydrophobic nature of Wnt, it is likely that the protein is interacting with other components or is even forming structures that would hide its lipid modification from the extracellular medium. In other systems it has been observed that specific proteins can bind and hide Wnt lipids, or that Wnt can travel on lipoprotein particles or exosomes, or even form micelles ^8,32^. Thus, accessory proteins of unknown identity might be of crucial interest to control biophysical parameters like the dynamics and the spread of the Wnt ligands in the tissue, as well as biochemical properties controlling the activity of the Wnt pathway. The combination of both effects might act as an extracellular regulation of Wnt activity to ensure proper signaling. Identifying these partners will be essential to understand the mechanisms of spreading and signaling in developing tissues. It is important to note that our results do not prove the existence of a concentration gradient of free Wnt in the tissue, either at short or long distances. We rather show that there are Wnt clusters distributed in an inhomogeneous manner [Figure 1a], and it is unclear how such a noisy distribution could be instructive for patterning of the developing tissue, even though complementary mechanisms could result in robust signaling from an imprecise distribution ^4,7^. Moreover, our observations do not rule out the possibility that a fraction of Wnt spreading is due to cytoneme- or filopodia-based mechanisms, even though we have not been able to detect such structures in *C. elegans* embryos.

Besides characterizing Wnt spreading in *C. elegans* embryos, we looked at the distribution of its receptor MOM-5/Fz. Beyond a possible genetic regulation that lower levels of Fz in the regions where Wnt concentration is high, we observed that the receptor itself forms clusters on the membrane of most of the embryonic cells and that these clusters partially colocalize with Wnt punctae. This result suggests a direct interaction between the ligand and its receptor and a possible role of one in the organization of the other. Besides, we found an overall concentration dependence of Fz with CWN-2 [Figure 3] and observed only partial colocalization with CWN-2. Even if it is likely that other ligands (CWN-1, MOM-2) could also be responsible for forming Fz clusters in which CWN-2 is absent. We tried to study the organization of Fz in dissociated embryonic cells under different Wnt conditions. However, decreasing or increasing Wnt concentration in the culture medium gives similar results. These two conditions lead to a small increase in Fz concentration on the membrane compared to the control, while the density of clusters was not affected [Figure 4 – Supp Fig. 1]. Similarly, these two conditions had the same effect on the dynamics of Fz, it led to an increase in Fz diffusion coefficient compared to control. It is likely that Wnt concentration not only affects Fz distribution and mobility in the membrane but is also responsible for changes in Fz expression ^33–36^, and on the recycling of Fz from the membrane through endocytosis ^37,38^. Consequently, it is difficult to draw definitive conclusions from this experiment.

If the role of Wnt ligands in establishing cellular polarity of fields of cells is controversial in some systems ^39^, evidence exists that Wnt ligands have a role in transmitting polarity information, either by localizing membrane components or orienting spindles ^18,24,25^. However, the mechanisms at play are still unclear due to the variety of distributions and modes of spreading that Wnt ligands exhibit in different systems ^1,4,6,11,24,40–42^. These various modes of spreading are likely to achieve context-dependent functions. For example, it is accepted that a diffusion-based morphogen gradient does not perform well over large distances because of the time needed to spread, but a concentration gradient can also be perturbed by the growth or reorganization of a tissue. These perturbations are not necessarily compatible with the functional integration of a signal by a target cell. A cytoneme-based mechanism has been proposed as an alternative mode of transport of the ligands. This model has its own time and spatial scales, but it also requires tight regulations to achieve a precise spatial patterning of tissues ^8,43,44^. In *C. elegans* embryos, cells divide every 20-30 minutes on average. In this short time window, Wnt target cells must integrate information and take subsequent decisions. We propose that the freely diffusing Wnt population could be a carrier of polarity information without the necessity to establish a stable concentration gradient. A concentration gradient that would anyway be likely perturbed by cell-cell contact rearrangements in the developing tissue. Ligands produced in the posterior half of the embryo can spread over a few minutes in the anterior half, where they can bind Fz. Our model necessitates the Fz receptor to diffuse in the plasma membrane, which we showed is the case. This transfer of polarity from the tissue scale to the cellular scale does not require an active mechanism of spreading or complex signaling regulations. It relies only on the diffusion properties of the ligand and its receptor, properties that are sufficient to achieve polarization on the timescales and distances relevant to *C. elegans* embryos, but that might not be the case in other systems of larger sizes. Besides, this simple mechanism only applies to polarity transfer and not to tissue patterning that relies on thresholds of Wnt activity.

To better understand the potential role of freely diffusing Wnt, it will be necessary to characterize how they spread in the extracellular space. In particular, it will be important to identify the potential interactors that promote the solubility of the Wnt ligands. It will also be important to study the potential asymmetric cellular distribution of Fz in living embryos and the direct interaction of the receptor with the ligand. This work would benefit from advances in microscopy techniques to achieve nanometer resolution in living tissues (to distinguish Fz on membranes of neighbouring cells) as well as advances in tagging endogenous proteins with organic fluorophores (to track at high framerate diffusing molecules). This information would help clarify the role of the freely diffusing population as well as the nature of the molecular organization of stable Wnt and Fz clusters on the plasma membrane.

## Material and Methods

### *C. elegans* animals

*C. elegans* animals were grown on Normal Growth Media (NGM) plates, fed with E. coli OP50 and cultured at 20°C for experiments. *mom-5endo::mKate2* (*vba10*) knock-in strain was generated using CRISPR methods ^16^, with insertion of the fluorescent protein at the C-terminus of *mom-5*. The sequence targeted by the guide RNA is TGTTGATCAGGTTAATATG<u>AGG</u> (PAM underlined). Homology arms (PCR products from genomic DNA) were inserted with Gibson assembly (New England Biolabs) into pDD287 (Addgene #70685), a plasmid that contains mKate2 fluorescent protein and a self-excising selection cassette. MOM-5 and the fluorescent protein are separated by a flexible linker (GASGASGAS). Candidate knock-ins were selected by drug treatment and phenotypic identification. Excision of the selection cassette was induced by heat-shock expression of Cre in L1/L2 larvae (4 hours at 34°C). Candidate knock-ins were checked with fluorescence imaging and PCR genotyping. The integrated array *phsp::cwn-2::mcherry* (*vbaIs29*) was generated by X-ray irradiation of an extrachromosomal array from ^45^.

### Imaging conditions and image analysis

Embryos were dissected from gravid adults in egg buffer and mounted on a 5% agarose pad between a glass slide and a coverslip. They were incubated at 20°C until observation. Fluorescence imaging was performed on two spinning disk confocal microscopes equipped with CSU-X1 spinning disk unit (Yokogawa) mounted on Nikon Ti eclipse stands: (1) 100x/1.49NA Nikon objective, EMCCD Andor iXon3 DU897, MicroManager software ^46^, (2) 100x/1.4 Nikon objective, EMCCD Photometrics Evolve (Teledyne), Metamorph software (Molecular devices). Images were acquired with excitation at 515nm and/or 561nm.

All image processing and data analysis were done with ImageJ and/or Python. Graphics were created with Python and Matplotlib library and assembled with Inkscape.

For the time evolution of Wnt strains fluorescence intensities, we acquired stack images (slice every 1 µm) and sum-projected the first 5 µm of the ventral view after background subtraction. Embryos were imaged every 20 minutes. Average fluorescence intensity is measured in a hand-drawn ROI corresponding to the embryo. This ROI is then separated into two halves (anterior and posterior) in which fluorescence intensity is measured. The polarity ratio is then defined as the posterior to the anterior intensity ratio. Profiles along the anteroposterior axis are measured along a line of 20µm at stages of interest. They are then normalized to their maximum length so that 0 corresponds to the anterior extremity and 1 to the posterior extremity.

Clusters of CWN-2::YFP or MOM-5::mKate2 (in embryos or single cells) are detected in profile manually created along the cell plasma membrane, either in the anterior half or the posterior half of the embryo. Clusters were thresholded automatically. The threshold value for each image was defined as follows: histograms of intensity in the profiles are computed, and the background (most frequent intensity values) is fitted with a gaussian curve to obtain a fluorescence intensity average value I_avg_ and a variance I_var_. Clusters are then detected along the profile as the pixels with an intensity higher than I_avg_ + 2*I_var_ using the *find_peaks* function from the *SciPy*.*signal* library ^47^.

Fluorescence Recovery After Photobleaching was performed on VBS322 strains. The anterior half of 4h old embryos were bleached with a 515nm laser (Metamorph with FRAP module). Fluorescence recovery was measured every 2 min for 16 min. CWN-2::YFP clusters were detected automatically with ImageJ plugin TrackMate (spot radius = 0.8µm, median filtering On, threshold = 20) ^26,48^. We counted the number of clusters over time in two compartments of the anterior half of 16 embryos and normalized it with the initial (pre-bleach) number of clusters. The average recovery in the compartment closest to the source was fitted with an exponential 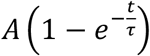 with A the maximum recovery and *τ* the characteristic recovery time.

### Fluorescence Correlation Spectroscopy

Fluorescence Correlation Spectroscopy (FCS) was performed on a Zeiss LSM780 confocal microscope equipped with a 40x/1.2NA Water immersion Zeiss objective. Samples were excited with a 515nm laser and acquisition was made in 525-700nm band on a GaAsP detector. The focal point was positioned in the volume between the egg shell and the tissue. Fluorescence intensity correlograms were analyzed with open-source graphical user interface PyCorrFit ^49^ to extract diffusion coefficients of MOM-2::YFP molecules. We fitted autocorrelograms with a model including triplet state and two-component three-dimensional free diffusion. The fast characteristic time component is associated with the photophysics of yellow fluorescent protein ^50^, while the longer characteristic time (about 1ms) corresponds to the diffusion process of the molecule. The latter was used to estimate the diffusion coefficient with the following formula: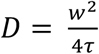, with 3 the waist of the confocal volume.

### *C. elegans* embryonic primary cell culture

Before *C. elegans* embryonic primary cell cultures, two strains, LP184 (*mom-5::YPet*) and VBS199 (*hsp::cwn-2::mCherry*), were grown in liquid culture ^51,52^. The liquid culture was started using three 65mm petri plate full of worms using the synchronization technique mentioned in ^53–55^. The embryonic cell isolations were then started with two tubes of 400µl precipitated synchronized egg-laying adults of both strains. Each tube was washed thrice with M9. VBS199 was heat shocked for 20 minutes at 37°C. The worms were then lysed using NaOH and bleach. Eggs were collected in one tube for each strain. Both tubes were washed thrice with egg buffer, and eggs were collected by centrifugation. Eggs and debris are separated using 30% sucrose solution in egg buffer followed by washing off sucrose. The pellets were dissolved in egg buffer. To digest the eggshell sample was treated with chitinase, and the reaction was stopped by Leibovitz’s L-15 medium supplementation. The eggs were collected by centrifugation, and the pellets were dissolved in fresh L-15. The samples were pipetted in three consecutive steps using a 100-1000μl micro-pipette to dissociate the eggs gently. Each step includes pipetting the samples seventy-five times, followed by centrifugation at 100g and collection of the dissociated cell containing supernatant. After this step, the sample was a heterogeneous mixture of dissociated single cells and small cell clusters along with undissociated eggs, hatched larvae and eggshells. The samples were centrifuged 900g, and the cloudy supernatants were removed (small debris). The pellet was dissolved in L-15 by pipetting the sample in and out ∼50 times and centrifuged at 100g. The supernatants were collected without disturbing the pellet as dissociated cells are present in the supernatant, and large debris are present in the pellet. Cell density was measured, and aliquots were made. The Wnt secretion inhibitor LGK974 (Cayman Chemical) was added if required (100nM). The LP184 cells were plated on acid-washed and peanut lectin-treated glass surface and incubated for 1hr15. The samples were washed three times with L-15 medium to remove nonattached cells. VBS199 cells were added if necessary.

### Single molecule tracking

MOM-5::YPET was imaged in single embryonic cells at a framerate of 50 ms with a spinning disk microscope. Movies (700 images) were analyzed with FiJi and its plugin TrackMate ^26,48^. The first 300 images were discarded to enter single molecule mode (low density of fluorescent molecules after photobleaching). After background subtraction, movies are batch processed to detect single molecules with a Laplacian of Gaussian detector. Trajectories were obtained with LAPtracker procedure. MSD curves were computed and analyzed with a Python algorithm that returns the diffusion coefficient D and anomalous exponent α after fitting with the function 452(7) = 427^)^. Average diffusion coefficients are estimated from MSD curves with an anomalous exponent between 0.9 and 1.1. Detailed protocol and scripts are available at https://github.com/remyeltorro/Batch-SPT/tree/main/notes.

### Simulations

Reaction diffusion simulations were performed using SmolDyn software ^28,29,56^. Embryos were modelled as a 2D array of 12 by 8 square cells (5µm width) contained in a impermeable box modelling the eggshell. Cells were separated by 50 nm. The rightmost column of cells is the source of 4000 Wnt ligands, which are secreted from these cells in the extracellular space. They diffuse in that space with a diffusion coefficient set to 2µm^2^/s. On the membrane of all cells, we placed 40 Fz molecules which can diffuse in the membrane with a diffusion coefficient of 0.1 µm^2^/s. Diffusion coefficient are set according to experimental results obtained from FCS and single molecule tracking. A diffusion limited reaction is set between Wnt ligands and Fz receptor. Upon formation this complex is not diffusing in the membrane, to allow for the localization of the interaction. Simulations configuration files and analysis procedures were created with a custom-made code written in Python, available on https://github.com/s2raf/WntFz.

---

| Name | Genotype | Reference |
| --- | --- | --- |
| VBS319 | *vbals40[mom-2::YFP]; itls44[PH::mCherry]* | 18 |
| VBS320 | *vbals39[cwn-1::YFP]; itls44[PH::mCherry]* | 18 |
| VBS322 | *vbals43[cwn-2::YFP]; itls44[PH::mCherry]* | 18 |
| VBS429 | *vbals43[cwn-2::YFP]; mom-5(vba10[mom-5::mKate2]) I* | This study |
| LP184 | *mom-5(cp31[mom-5::YPET]) I* | Heppert et al., 2018 |
| VBS488 | *mom-5(cp31[mom-5::YPET]) I; itls44[PH::mCherry]* | This study |
| VBS199 | *vbals29 [phsp::cwn-2::mCherry]* | This study |
| VBS471 | *mom-5(gk812) I/hT2 [bli-4(e937) let-?(q782) qIs48] (I;III); vbals43[cwn-2::YFP]* | This study |

**Figure 1-figure supplement 1.**
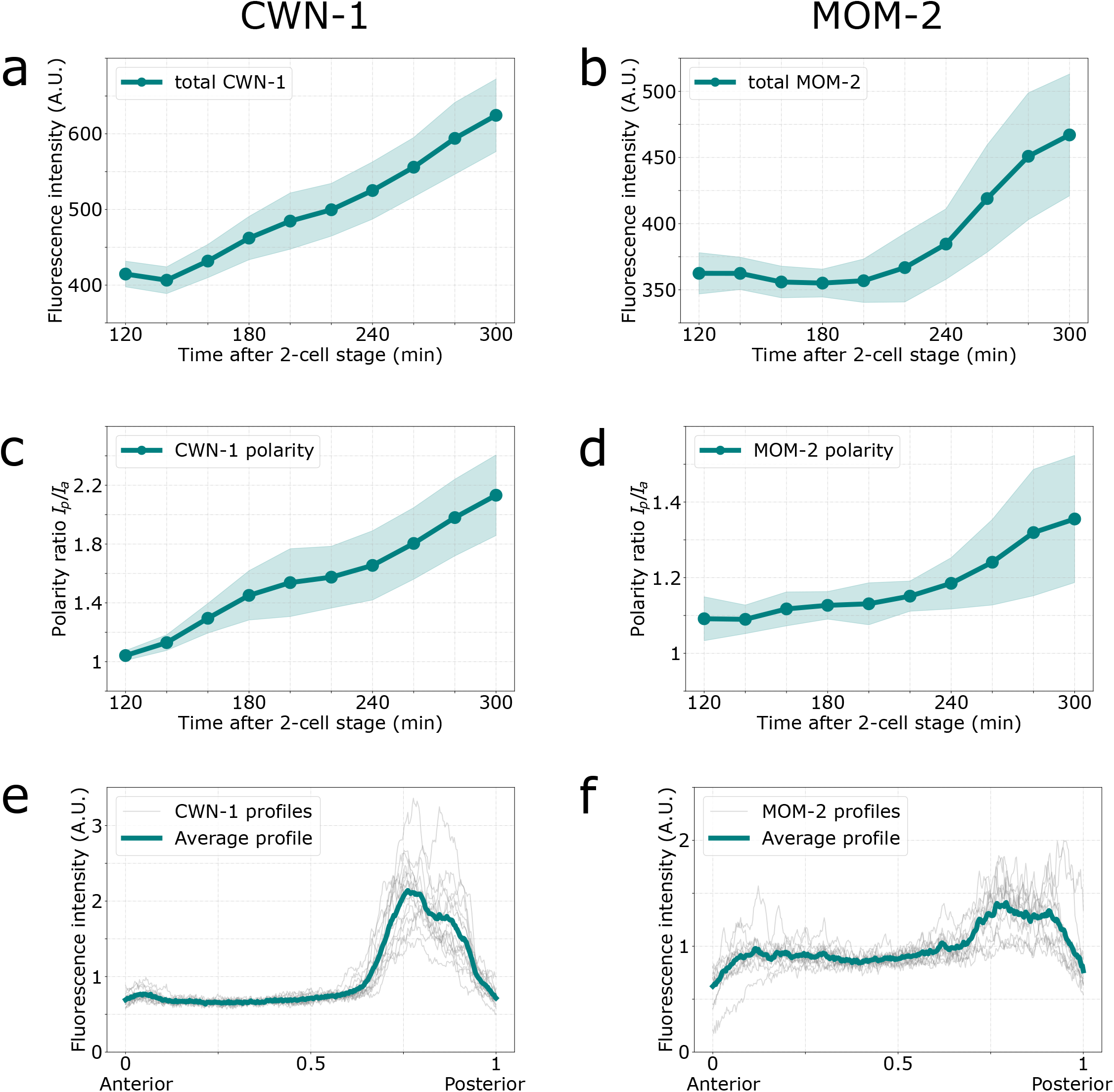
Time evolutions and distributions of tagged Wnt ligands. a & b- Time evolution of the total fluorescence intensities of CWN-1::YFP and MOM-2::YFP expressing embryos. Acquisitions start 2 hours after the 2-cell stage. c & d- Time evolutions of the polarity ratios of CWN-1::YFP and MOM-2::YFP expressing embryos. Acquisitions start 2 hours after 2- cell stage. e & f- Fluorescence intensity profiles along the antero-posterior axis of CWN-1::YFP and MOM-2::YFP expressing embryos 4 hours after 2-cell stage.

**Figure 4-supplement figure 1.**
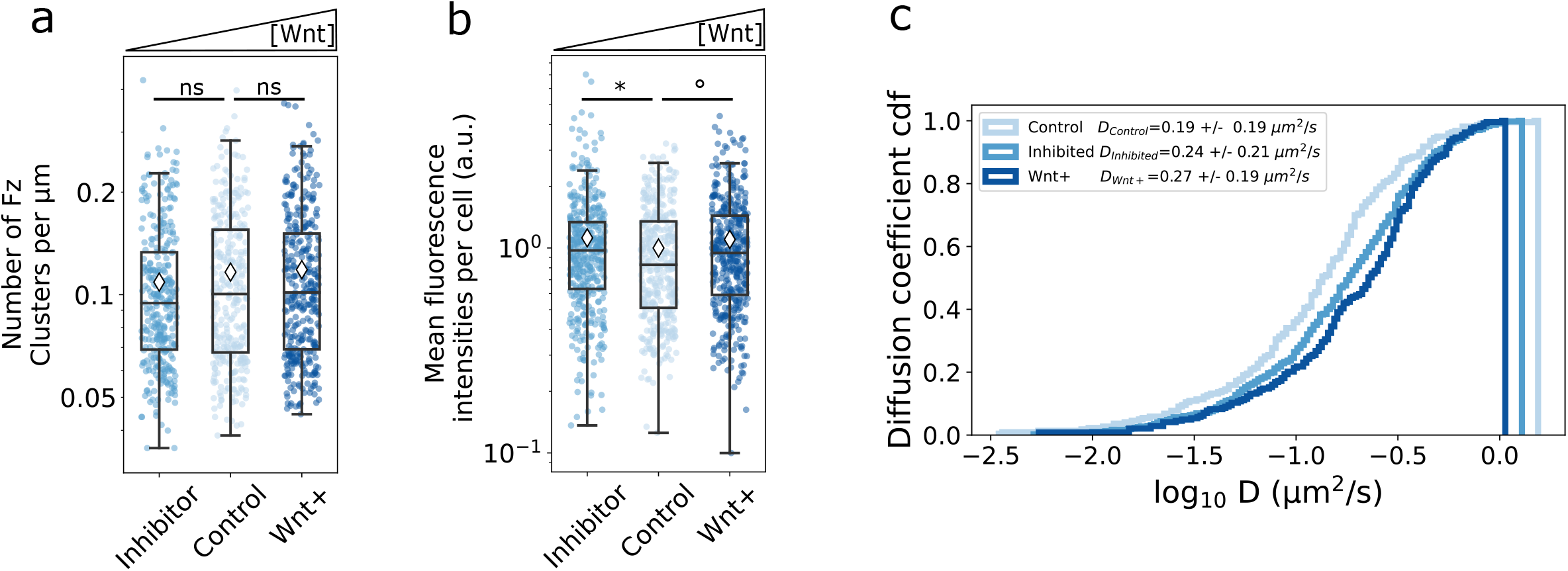
Effect of Wnt concentration on the distribution of Fz on the plasma membrane of isolated embryonic cells in culture. Wnt concentration in the medium was reduced compared to the *control* condition by treatment with a Wnt secretion inhibitor (LGK974 – Cayman Chemical)^57^, *inhibitor* condition. Wnt concentration was increased by co-culturing *cp31* cells (MOM-5::YPET) with *vbaIs29* (*phsp::cwn-2::mCherry*) cells that overexpress CWN-2 after heat shock. *Wnt+* condition. a- Number of clusters in single cells in different conditions. b- Mean fluorescence intensity (MOM-5::YPET) on the plasma membrane of single cells in different conditions.(* p= 0.03; ° p=0.02, Mann-Whitney tests) c- Cumulative distribution functions of the MOM-5 diffusion coefficient in different conditions.

**Movie 1.**
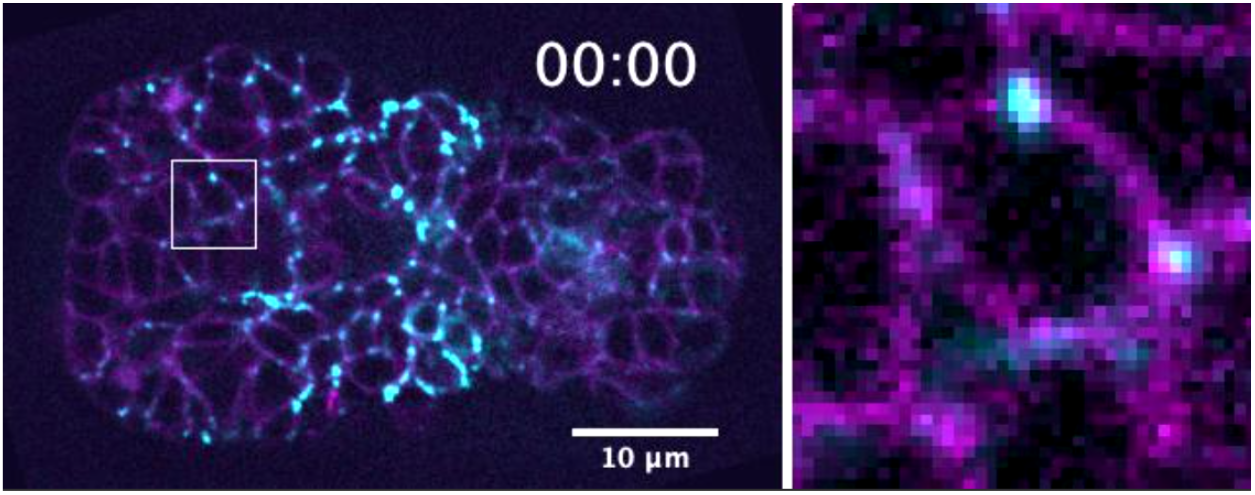
Time evolution of Wnt/CWN-2::YFP (cyan) and PH::mCherry (membrane marker in magenta) in a 5h old *C. elegans* embryo. Right: zoom of the cells in white square.

**Movie 2.**
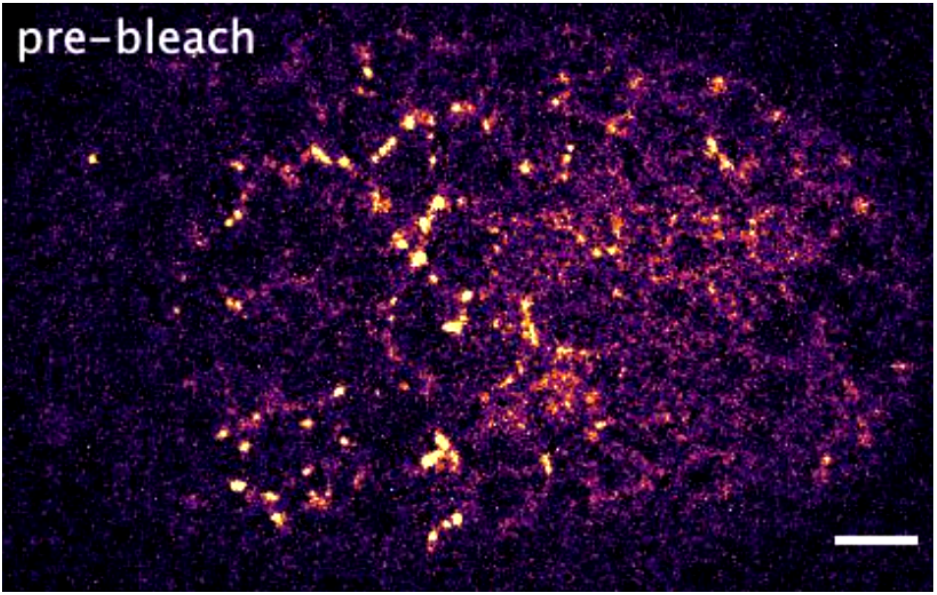
Fluorescence Recovery After Photobleaching in a 4h old embryo of CWN-2::YFP. The anterior half of the embryo was bleached at time 0. Scale bar is 5µm.

**Movie 3.**
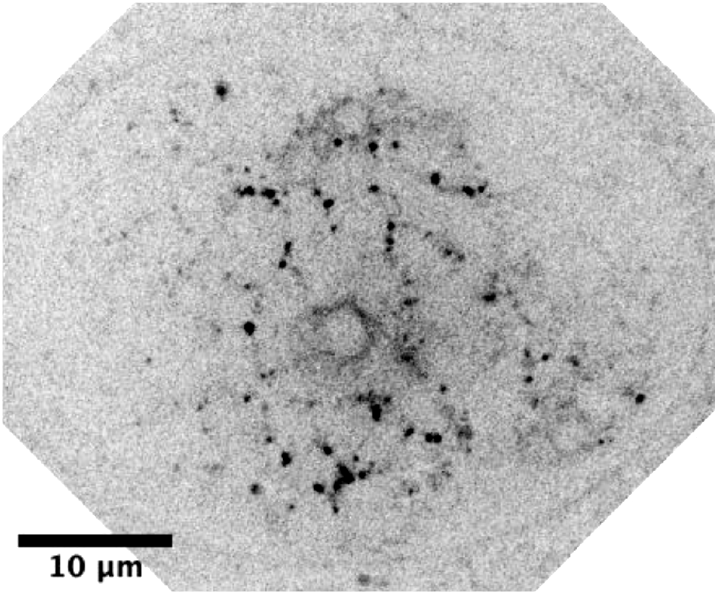
Stack images of an embryo expressing CWN-2::YFP in the background of *mom-5* mutation (*gk812*). One slice every 0.5 µm.

**Movie 4.**
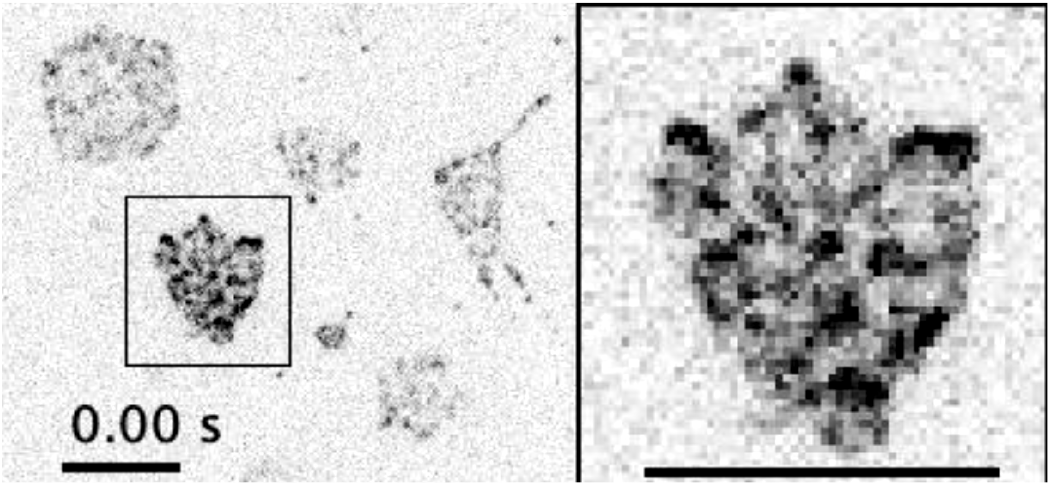
Single molecule movie of MOM-5::YPET. Right, zoom in on the cell in the black square. In the first images, the signal is too dense to distinguish single molecules. After a bleaching phase, the density of observable molecules is adapted for single-molecule tracking. Scale bar is 5 µm.

**Movie 5.**
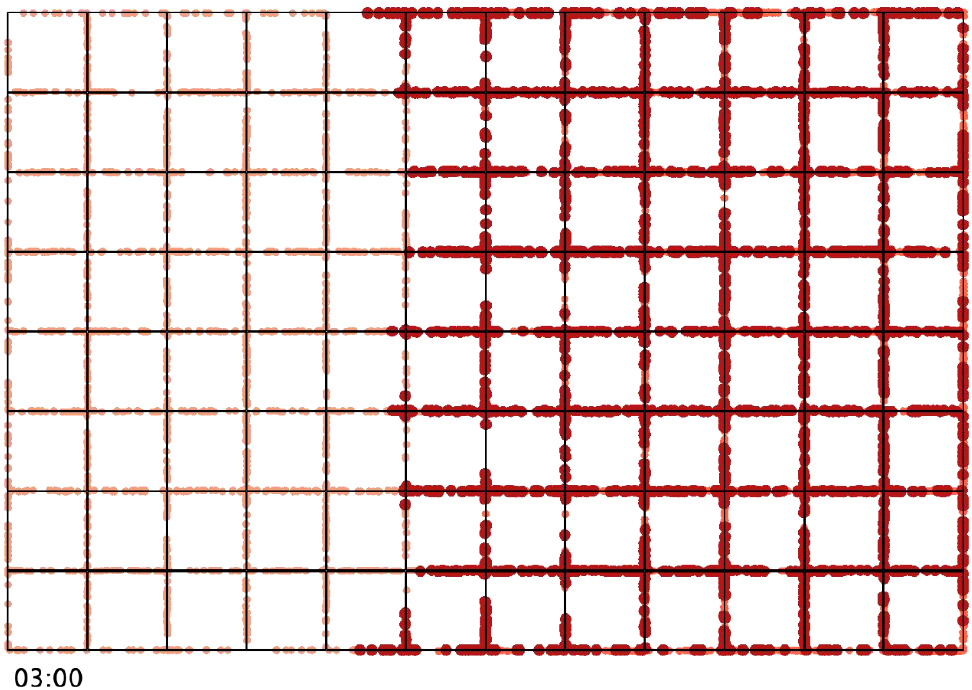
Movie from a simulation of diffusion of Wnt ligands in the extracellular space and binding with a membrane receptor. Simulated time in min:sec. Geometry: 96 cells, 4 µm each, separated by 50 nm. Diffusion coefficient of Wnt is 2 µm^2^/s, diffusion coefficient of Fz is 0.1 µm^2^/s. The complex formed by Wnt and Fz is immobile on the membrane to mark the location of their formation on the plasma membrane. One can observe a wave of formation of the complex on the cells, from the source of Wnt toward the anterior part of the embryo.

## Acknowledgements

This work was funded by grants from the Agence Nationale de la Recherche (ANR-14-CE11-0001 and ANR-11-LABX-0054) to V.B. and P.-F.L. Pritha Pai was funded by a grant from Fondation Pour la Recherche Médicale (FRM, grant FDT201904008378). Imaging was performed on a PiCSL-FBI core facility (IBDM, AMU-Marseille) supported by the French National Research Agency through the ‘Investments for the Future’ program (France-BioImaging, ANR-10INBS-04). Some strains were provided by the Caenorhabditis Genetics Center, which is funded by NIH Office of Research Infrastructure Programs (P40 OD010440). The authors thank all the members of Lenne and Bertrand labs and Pierre Mangeol for fruitful discussion and reading of the manuscript. We also thank O. Theodoly for conceptual and technical support. P.R. thanks all the interns he supervised who helped along the way.

## Author contributions

<u>Pierre Recouvreux</u>, Conceptualization, Formal analysis, Funding acquisition, Investigation, Methodology, Software, Resources, Visualization, Supervision, Writing - original draft; <u>Pritha Pai</u>, Conceptualization, Formal analysis, Investigation, Methodology, Resources, Writing - original draft; <u>Rémy Torro</u>, Formal analysis, Investigation, Methodology, Software; <u>Mónika Ludányi</u>, Investigation, Resources; <u>Pauline Mélénec</u>, Investigation, Resources; <u>Mariem Boughzala</u>, Investigation, Resources; <u>Vincent Bertrand</u>, Conceptualization, Formal analysis, Funding acquisition, Investigation, Methodology, Writing original draft; Pierre-François Lenne, Conceptualization, Formal analysis, Funding acquisition, Investigation, Methodology, Supervision, Writing original draft.

## BIBLIOGRAPHY

1. Coudreuse, D.Y.M., Roël, G., Betist, M.C., Destrée, O., and Korswagen, H.C. (2006). Wnt Gradient Formation Requires Retromer Function in Wnt-Producing Cells. Science 312, 921–924. 10.1126/science.1124856.

2. Gregor, T., Wieschaus, E.F., McGregor, A.P., Bialek, W., and Tank, D.W. (2007). Stability and Nuclear Dynamics of the Bicoid Morphogen Gradient. Cell 130, 141–152. 10.1016/j.cell.2007.05.026.

3. Müller, P., and Schier, A.F. (2011). Extracellular Movement of Signaling Molecules. Dev. Cell 21, 145–158. 10.1016/j.devcel.2011.06.001.

4. Pani, A.M., and Goldstein, B. (2018). Direct visualization of a native Wnt in vivo reveals that a long-range Wnt gradient forms by extracellular dispersal. eLife 7, e38325. 10.7554/eLife.38325.

5. Yu, S.R., Burkhardt, M., Nowak, M., Ries, J., Petrášek, Z., Scholpp, S., Schwille, P., and Brand, M. (2009). Fgf8 morphogen gradient forms by a source-sink mechanism with freely diffusing molecules. Nature 461, 533–536. 10.1038/nature08391.

6. Kicheva, A., Pantazis, P., Bollenbach, T., Kalaidzidis, Y., Bittig, T., Jülicher, F., and González-Gaitán, M. (2007). Kinetics of Morphogen Gradient Formation. Science 315, 521–525. 10.1126/science.1135774.

7. Muller, P., Rogers, K.W., Yu, S.R., Brand, M., and Schier, A.F. (2013). Morphogen transport. Development 140, 1621–1638. 10.1242/dev.083519.

8. Stapornwongkul, K.S., and Vincent, J.-P. (2021). Generation of extracellular morphogen gradients: the case for diffusion. Nat. Rev. Genet. 22, 393–411. 10.1038/s41576-021-00342-y.

9. Abu-Arish, A., Porcher, A., Czerwonka, A., Dostatni, N., and Fradin, C. (2010). High Mobility of Bicoid Captured by Fluorescence Correlation Spectroscopy: Implication for the Rapid Establishment of Its Gradient. Biophys. J. 99, L33–L35. 10.1016/j.bpj.2010.05.031.

10. Petersen, C.P., and Reddien, P.W. (2009). Wnt Signaling and the Polarity of the Primary Body Axis. Cell 139, 1056–1068. 10.1016/j.cell.2009.11.035.

11. Mii, Y., Nakazato, K., Pack, C.-G., Ikeda, T., Sako, Y., Mochizuki, A., Taira, M., and Takada, S. (2021). Quantitative analyses reveal extracellular dynamics of Wnt ligands in Xenopus embryos. eLife 10, e55108. 10.7554/eLife.55108.

12. Farin, H.F., Jordens, I., Mosa, M.H., Basak, O., Korving, J., Tauriello, D.V.F., de Punder, K., Angers, S., Peters, P.J., Maurice, M.M., et al. (2016). Visualization of a short-range Wnt gradient in the intestinal stem-cell niche. Nature 530, 340–343. 10.1038/nature16937.

13. Bertrand, V. (2016). β-catenin-driven binary cell fate decisions in animal development. WIREs Dev. Biol. 5, 377–388. 10.1002/wdev.228.

14. Chen, C.-K., and Pan, C.-L. (2022). Cell polarity control by Wnt morphogens. Dev. Biol. 487, 34–41. 10.1016/j.ydbio.2022.04.007.

15. Sawa, H. (2012). Control of Cell Polarity and Asymmetric Division in C. elegans. In Current Topics in Developmental Biology (Elsevier), pp. 55–76. 10.1016/B978-0-12-394592-1.00003-X.

16. Dickinson, D.J., Pani, A.M., Heppert, J.K., Higgins, C.D., and Goldstein, B. (2015). Streamlined Genome Engineering with a Self-Excising Drug Selection Cassette. Genetics 200, 1035–1049. 10.1534/genetics.115.178335.

17. Heppert, J.K., Pani, A.M., Roberts, A.M., Dickinson, D.J., and Goldstein, B. (2018). A CRISPR Tagging-Based Screen Reveals Localized Players in Wnt-Directed Asymmetric Cell Division. Genetics 208, 1147–1164. 10.1534/genetics.117.300487.

18. Kaur, S., Mélénec, P., Murgan, S., Bordet, G., Recouvreux, P., Lenne, P.-F., and Bertrand, V. (2020). Wnt ligands regulate the asymmetric divisions of neuronal progenitors in C. elegans embryos. Development 147, dev183186. 10.1242/dev.183186.

19. Sawa, H., and Korswagen, H.C. (2013). Wnt signaling in C. elegans. WormBook, 1–30. 10.1895/wormbook.1.7.2.

20. Harterink, M., Kim, D. hyun, Middelkoop, T.C., Doan, T.D., Oudenaarden, A. van, and Korswagen, H.C. (2011). Neuroblast migration along the anteroposterior axis of C. elegans is controlled by opposing gradients of Wnts and a secreted Frizzled-related protein. Development 138, 2915–2924. 10.1242/dev.064733.

21. Zacharias, A.L., Walton, T., Preston, E., and Murray, J.I. (2015). Quantitative Differences in Nuclear β-catenin and TCF Pattern Embryonic Cells in C. elegans. PLOS Genet. 11, e1005585. 10.1371/journal.pgen.1005585.

22. Takada, R., Mii, Y., Krayukhina, E., Maruyama, Y., Mio, K., Sasaki, Y., Shinkawa, T., Pack, C.-G., Sako, Y., Sato, C., et al. (2018). Assembly of protein complexes restricts diffusion of Wnt3a proteins. Commun. Biol. 1, 1–14. 10.1038/s42003-018-0172-x.

23. Park, F.D., Tenlen, J.R., and Priess, J.R. (2004). C. elegans MOM-5/Frizzled Functions in MOM-2/Wnt-Independent Cell Polarity and Is Localized Asymmetrically prior to Cell Division. Curr. Biol. 14, 2252–2258. 10.1016/j.cub.2004.12.019.

24. Goldstein, B., Takeshita, H., Mizumoto, K., and Sawa, H. (2006). Wnt Signals Can Function as Positional Cues in Establishing Cell Polarity. Dev. Cell 10, 391–396. 10.1016/j.devcel.2005.12.016.

25. Habib, S.J., Chen, B.-C., Tsai, F.-C., Anastassiadis, K., Meyer, T., Betzig, E., and Nusse, R. (2013). A Localized Wnt Signal Orients Asymmetric Stem Cell Division in Vitro. Science 339, 1445–1448. 10.1126/science.1231077.

26. Tinevez, J.-Y., Perry, N., Schindelin, J., Hoopes, G.M., Reynolds, G.D., Laplantine, E., Bednarek, S.Y., Shorte, S.L., and Eliceiri, K.W. (2017). TrackMate: An open and extensible platform for single-particle tracking. Methods 115, 80–90. 10.1016/j.ymeth.2016.09.016.

27. Michalet, X. (2010). Mean square displacement analysis of single-particle trajectories with localization error: Brownian motion in an isotropic medium. Phys. Rev. E 82, 041914. 10.1103/PhysRevE.82.041914.

28. Andrews, S.S. (2012). Spatial and Stochastic Cellular Modeling with the Smoldyn Simulator. In Bacterial Molecular Networks Methods in Molecular Biology., J. van Helden, A. Toussaint, and D. Thieffry, eds. (Springer New York), pp. 519–542. 10.1007/978-1-61779-361-5_26.

29. Andrews, S.S., and Bray, D. (2004). Stochastic simulation of chemical reactions with spatial resolution and single molecule detail. Phys. Biol. 1, 137–151. 10.1088/1478-3967/1/3/001.

30. Mii, Y., and Taira, M. (2009). Secreted Frizzled-related proteins enhance the diffusion of Wnt ligands and expand their signalling range. Development 136, 4083–4088. 10.1242/dev.032524.

31. Mii, Y., Yamamoto, T., Takada, R., Mizumoto, S., Matsuyama, M., Yamada, S., Takada, S., and Taira, M. (2017). Roles of two types of heparan sulfate clusters in Wnt distribution and signaling in Xenopus. Nat. Commun. 8, 1973. 10.1038/s41467-017-02076-0.

32. Langton, P.F., Kakugawa, S., and Vincent, J.-P. (2016). Making, Exporting, and Modulating Wnts. Trends Cell Biol. 26, 756–765. 10.1016/j.tcb.2016.05.011.

33. Cadigan, K.M., Fish, M.P., Rulifson, E.J., and Nusse, R. (1998). Wingless Repression of Drosophila frizzled 2 Expression Shapes the Wingless Morphogen Gradient in the Wing. Cell 93, 767–777. 10.1016/S0092-8674(00)81438-5.

34. Muller, H.-A., Samanta, R., and Wieschaus, E. (1999). Wingless signaling in Drosophila embryo: zygotic requirements and the role of the frizzled genes. Development, 577–586.

35. Sato, A., Kojima, T., Ui-Tei, K., Miyata, Y., and Saigo, K. (1999). Dfrizzled-3, a new Drosophila Wnt receptor, acting as an attenuator of Wingless signaling in wingless hypomorphic mutants. Development 126, 4421–4430. 10.1242/dev.126.20.4421.

36. Willert, J., Epping, M., Pollack, J.R., Brown, P.O., and Nusse, R. (2002). A transcriptional response to Wnt protein in human embryonic carcinoma cells. BMC Dev. Biol., 7.

37. Blitzer, J.T., and Nusse, R. (2006). A critical role for endocytosis in Wnt signaling. BMC Cell Biol. 7, 28. 10.1186/1471-2121-7-28.

38. Brunt, L., and Scholpp, S. (2018). The function of endocytosis in Wnt signaling. Cell. Mol. Life Sci. 75, 785–795. 10.1007/s00018-017-2654-2.

39. Yu, J.J.S., Maugarny-Calès, A., Pelletier, S., Alexandre, C., Bellaiche, Y., Vincent, J.-P., and McGough, I.J. (2020). Frizzled-Dependent Planar Cell Polarity without Secreted Wnt Ligands. Dev. Cell 54, 583-592.e5. 10.1016/j.devcel.2020.08.004.

40. Alexandre, C., Baena-Lopez, A., and Vincent, J.-P. (2014). Patterning and growth control by membrane-tethered Wingless. Nature 202, 11. 10.1038/nature12879.

41. Chaudhary, V., Hingole, S., Frei, J., Port, F., Strutt, D., and Boutros, M. (2019). Robust Wnt signaling is maintained by a Wg protein gradient and Fz2 receptor activity in the developing Drosophila wing. Development 146, dev174789. 10.1242/dev.174789.

42. Lu, M., and Mizumoto, K. (2019). Gradient-independent Wnt signaling instructs asymmetric neurite pruning in C. elegans. eLife 8, e50583. 10.7554/eLife.50583.

43. Bressloff, P.C., and Kim, H. (2019). Search-and-capture model of cytoneme-mediated morphogen gradient formation. Phys. Rev. E 99, 052401. 10.1103/PhysRevE.99.052401.

44. Rosenbauer, J., Zhang, C., Mattes, B., Reinartz, I., Wedgwood, K., Schindler, S., Sinner, C., Scholpp, S., and Schug, A. (2020). Modeling of Wnt-mediated tissue patterning in vertebrate embryogenesis. PLOS Comput. Biol. 16, e1007417. 10.1371/journal.pcbi.1007417.

45. Jensen, M., Hoerndli, F.J., Brockie, P.J., Wang, R., Johnson, E., Maxfield, D., Francis, M.M., Madsen, D.M., and Maricq, A.V. (2012). Wnt Signaling Regulates Acetylcholine Receptor Translocation and Synaptic Plasticity in the Adult Nervous System. Cell 149, 173–187. 10.1016/j.cell.2011.12.038.

46. Edelstein, A., Amodaj, N., Hoover, K., Vale, R., and Stuurman, N. (2010). Computer Control of Microscopes Using µManager. Curr. Protoc. Mol. Biol. 92, 14.20.1-14.20.17. 10.1002/0471142727.mb1420s92.

47. Virtanen, P., Gommers, R., Oliphant, T.E., Haberland, M., Reddy, T., Cournapeau, D., Burovski, E., Peterson, P., Weckesser, W., Bright, J., et al. (2020). SciPy 1.0: fundamental algorithms for scientific computing in Python. Nat. Methods 17, 261–272. 10.1038/s41592-019-0686-2.

48. Ershov, D., Phan, M.-S., Pylvänäinen, J.W., Rigaud, S.U., Le Blanc, L., Charles-Orszag, A., Conway, J.R.W., Laine, R.F., Roy, N.H., Bonazzi, D., et al. (2022). TrackMate 7: integrating state-of-the-art segmentation algorithms into tracking pipelines. Nat. Methods 19, 829–832. 10.1038/s41592-022-01507-1.

49. Müller, P., Schwille, P., and Weidemann, T. (2014). PyCorrFit— generic data evaluation for fluorescence correlation spectroscopy. Bioinformatics 30, 2532–2533. 10.1093/bioinformatics/btu328.

50. Schwille, P., Kummer, S., Heikal, A.A., Moerner, W.E., and Webb, W.W. (2000). Fluorescence correlation spectroscopy reveals fast optical excitation-driven intramolecular dynamics of yellow fluorescent proteins. Proc. Natl. Acad. Sci. 97, 151–156. 10.1073/pnas.97.1.151.

51. Stiernagle, T. (2006). Maintenance of C. elegans. WormBook. 10.1895/wormbook.1.101.1.

52. Zhang, S., and Kuhn, J.R. (2013). Cell isolation and culture. WormBook, 1–39. 10.1895/wormbook.1.157.1.

53. Christensen, M., Estevez, A., Yin, X., Fox, R., Morrison, R., McDonnell, M., Gleason, C., Miller, D.M., and Strange, K. (2002). A Primary Culture System for Functional Analysis of C. elegans Neurons and Muscle Cells. Neuron 33, 503–514. 10.1016/S0896-6273(02)00591-3.

54. Sangaletti, R., and Bianchi, L. (2013). A Method for Culturing Embryonic C. elegans Cells. JoVE J. Vis. Exp., e50649. 10.3791/50649.

55. Strange, K., Christensen, M., and Morrison, R. (2007). Primary culture of Caenorhabditis elegans developing embryo cells for electrophysiological, cell biological and molecular studies. Nat. Protoc. 2, 1003–1012. 10.1038/nprot.2007.143.

56. Andrews, S.S., Addy, N.J., Brent, R., and Arkin, A.P. (2010). Detailed Simulations of Cell Biology with Smoldyn 2.1. PLOS Comput. Biol. 6, e1000705. 10.1371/journal.pcbi.1000705.

57. Liu, J., Pan, S., Hsieh, M.H., Ng, N., Sun, F., Wang, T., Kasibhatla, S., Schuller, A.G., Li, A.G., Cheng, D., et al. (2013). Targeting Wnt-driven cancer through the inhibition of Porcupine by LGK974. Proc. Natl. Acad. Sci. 110, 20224–20229. 10.1073/pnas.1314239110.

